# Extraocular Electrical Stimulation Activates Retinal Ganglion Cells In Vivo

**DOI:** 10.64898/2026.09.10.750688

**Authors:** Nicholas Householder, Omid Sharafi, Anahit Simonyan, Steven T. Walston, Gianluca Lazzi, Michael Bienkowski, Kimberly K. Gokoffski

**Affiliations:** Texas Tech University Health Science Center School of Medicine, Lubbock, Texas; Department of Electrical and Computer Engineering, Viterbi School of Engineering, University of Southern California, Los Angeles, California; Institute for Technology and Medical Systems (ITEMS), Keck School of Medicine, University of Southern California, Los Angeles, California; Department of Ophthalmology, Keck School of Medicine, USC Roski Eye Institute, University of Southern California, Los Angeles, California; Stevens Neuroimaging and Informatics Institute, Keck School of Medicine, University of Southern California, Los Angeles, California, United States of America; Zilkha Neurogenetic Institute, Keck School of Medicine, University of Southern California, Los Angeles, California, United States of America; Department of Ophthalmology and Vision Science, University of California, Irvine, Irvine, California

**Keywords:** Neurostimulation, Neuromodulation, Alternating Current Stimulation, Two-Photon Microscopy, Retinal Ganglion Cells

## Abstract

**Objective:** Here, we directly demonstrate that extraocular electrical stimulation can reliably activate retinal ganglion cells (RGCs) in vivo and systematically identify optimal stimulation waveforms that maximize RGC activation at tolerable amplitudes.

**Approach:** Using transpupillary two-photon calcium imaging in Thy1-GCaMP6f rats, we directly visualized RGC activation during extraocular electrical stimulation with single-cell resolution. We tested symmetric (SCB 1:1) and asymmetric (ACB 1:4) charge-balanced waveforms across frequencies ranging from 20 to 5,000 Hz and amplitudes from 1 to 300 µA, correlating cellular calcium responses with behavioral outcomes during awake stimulation.

**Main Results:** ACB 1:4 stimulation at lower frequencies (20–50 Hz) robustly and reliably activates RGC somas in vivo, producing larger calcium responses at lower amplitudes than SCB 1:1. At comparable amplitudes, ACB stimulation generated 1.8-fold greater calcium responses. In contrast, SCB stimulation required higher amplitudes that exceeded animal tolerance before reliable RGC activation could be achieved.

**Significance:** These findings provide direct evidence that extraocular electrodes can reliably activate RGCs in vivo. Notably, stimulation parameters previously associated with full-length optic nerve regeneration were found to be minimally effective at activating RGCs in vivo, suggesting increased gains could be had with newer approaches. The results further demonstrate that waveform asymmetry improves the efficiency of optic nerve stimulation by engaging RGC somas at lower, more tolerable, amplitudes. By combining two-photon imaging with behavioral tolerance testing, this work defines a practical therapeutic window for extraocular stimulation of the eye and establishes asymmetric charge-balanced waveforms as a more clinically translatable strategy for visual pathway neuromodulation.

## Introduction

Electric field (EF) stimulation of the visual pathway has emerged as a promising therapeutic approach for improving and restoring visual function in neuro-ophthalmic diseases such as amblyopia, glaucoma, optic neuritis, and traumatic optic nerve injury [1–3]. EF stimulation has been shown to elicit a wide range of desirable cellular responses, including axon growth, enhanced neuroplasticity, and improved neuronal survival [4–6]. In clinical studies, EF therapies have demonstrated favorable safety profiles, with tolerability and a low risk of serious adverse events [7–12].

Despite this promise, clinical translation of EF stimulation remains limited, in part, because stimulation strategies developed in vitro do not necessarily translate to effective and practical in vivo applications where electrode placement and stimulation constraints differ substantially [13].

Moreover, to be effective, many electro-regenerative strategies rely on placement of electrodes in direct contact with the retina (e.g. epiretinal or subretinal implants) inside the eye. [14–15] These electrodes cannot be routinely accessed when troubleshooting, revision, or upgrades are needed. Extraocular electrode configurations are of particular interest because they are more easily accessed when required. However, whether such configurations can reliably activate retinal ganglion cells (RGCs) in vivo across tolerable parameter ranges has not yet been directly established.

Here, we address this gap through two advances. First, we directly demonstrate that EF stimulation delivered through extra-ocular electrodes can reliably activate RGCs in vivo, establishing a practical method to engage the visual pathway. Second, we use this paradigm to systematically determine stimulation waveforms that most effectively drive RGC activation. By linking cellular activation measurements with behavioral tolerance constraints, this work defines a practical operating range for extra-ocular visual pathway stimulation and supports asymmetric charge-balanced stimulation as a more clinically viable strategy for visual pathway neuromodulation.

## Materials and Methods

### Electrode Installation

All animal procedures complied with the ARRIVE guidelines and the ARVO Statement for the Use of Animals in Ophthalmic and Vision Research, and were approved by the Institutional Animal Care and Use Committee at the University of Southern California [16] Briefly, adult (>300g) LE-Tg(Thy1-GCaMP6f)7 rats (830, Rat Resource & Research Center, University of Missouri, Columbia, MO, USA) were used [17]. A ‘J-shaped’ platinum source electrode (250 μm diameter; P1 Technologies, Boerne, TX) with 5 mm of its insulation coating stripped was routed subcutaneously from a midline scalp incision into the orbital cavity as seen in *figure 1a*. The electrode was maneuvered beneath the lateral rectus muscle and gently looped around the left optic nerve at the base of the globe to ensure stable contact without compressing the nerve. For feasibility testing, we utilized a straight needle made of platinum as reference electrode (250 μm diameter; P1 Technologies, Boerne, TX); this measures 16 mm in length, with 1 mm of insulation removed at the tip, and was implanted stereotactically into the contralateral optic tract using the following coordinates: 2.43 mm mediolateral (M-L), -1.00 mm anteroposterior (A-P), and -9.65 mm dorsoventral (D-V) at a 10° insertion angle (Stoelting 51900; Wood Dale, IL; *figure 1a*). The same electrodes, electrode positions, and surgical implantation procedures were performed as previously described [4].

**Figure 1:**
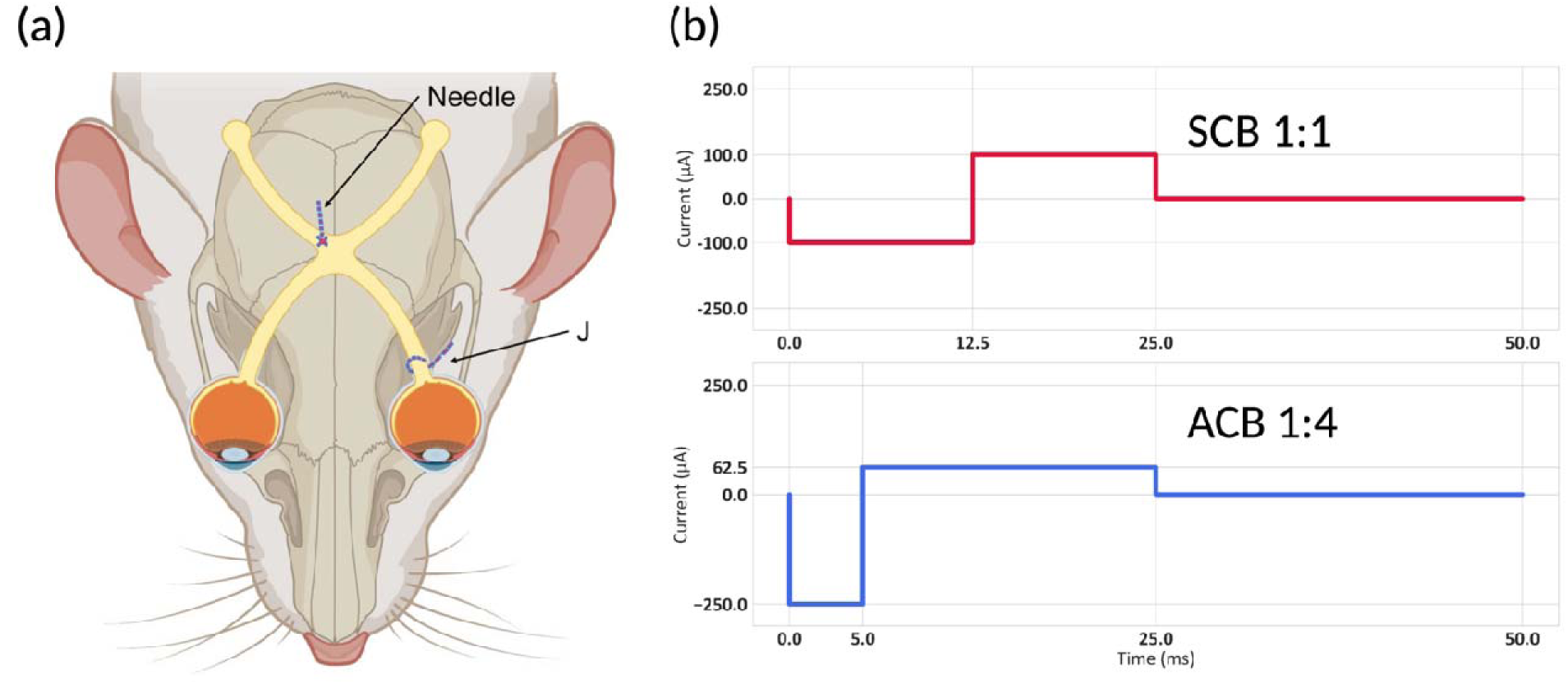
Schematic of electrode position and stimulation waveforms. (a) Schematic of the electrode position. (b) Diagram of SCB 1:1 and ACB 1:4 waveforms (20Hz).

### Electrical Stimulation Delivery and Parameter Design

Electrical stimulation was delivered using a constant-current stimulus isolator (Multi-Channel Systems STG series) connected to the two implanted electrodes via leads. Stimuli consist of biphasic alternating current (AC) waveforms applied in a charge-balanced manner to avoid net DC offset, which is known to cause cellular damage [18]. A symmetric charge-balanced (SCB) 1:1 waveform was compared with an ACB 1:4 waveform (figure 1b), as in [4].

Stimulation was applied at frequencies of 20, 50, 100, 500, 1000, and 5000 Hz (cycles per second) to probe frequency-dependent effects [19]. Each stimulus cycle was a continuous biphasic waveform, as previously described, with either a 1:1 or 1:4 ratio, 50% duty cycle (the percentage of time the stimulation is on, mathematically defined as *D = (t_ON_* ⍰ *T*) ×100%), and repeated at predetermined frequencies. The current amplitude was varied across trials from 1 to 300 µA to assess dose-dependent responses. The duty cycle was maintained at 50% for both the ACB and SCB waveforms, allowing for critical analysis of frequency-dependent effects without altering the total stimulation phase duration. To avoid cumulative effects, charge buildup, or order bias, stimulation frequencies and waveforms were randomly ordered and then circularly rotated, ensuring that each condition occurred in every sequence position, with a minimum of 3 min rest between new frequency sets. While frequency and waveform order were randomized, the amplitude was increased monotonically for each frequency and waveform to prevent delivering high-intensity stimulation early in the protocol, which could compromise animal safety, induce early motion artifacts, shift the imaging plane, and reduce the validity of subsequent trials.

The starting amplitudes were intentionally kept low to identify levels that could be delivered without causing movement; consequently, informative responses tended to emerge at higher amplitudes later in the sequence. To minimize carryover between trials, we ensured sufficient inter-stimulation rest to allow calcium-dependent fluorescence to return fully to baseline, as shown in figure 3. This precaution accounted for the known delay in GCaMP signal decay. It confirmed that responses from one trial did not influence those in the subsequent trial, thereby preserving the integrity of the data. Table 1 summarizes the key stimulation parameters used in this study, including the waveform shape, phase duration, frequency, and current amplitude.

**Table 1:** Electrical stimulation parameters. The ACB waveform delivered the cathodic phase at four times the amplitude for one-quarter of the duration of the opposite phase, achieving charge balance while maintaining an asymmetric pulse shape. The phase duration scales inversely with the frequency.

| Waveform | Phase Ratio<br>(cathodic:<br>anodic) | Frequency<br>Tested | Phase Duration per Cycle | Current<br>Amplitude |
| --- | --- | --- | --- | --- |
| **Symmetric (SCB 1:1)** | 1:1<br>(equal<br>amplitude and<br>duration) | 20, 50, 100,<br>500, 1000,<br>5000 Hz | 20Hz: 12.5 ms cathodic + 12.5 ms<br>anodic (DC = 50%)<br>50Hz: 5 ms cathodic + 5 ms anodic (DC<br>= 50%)<br>100Hz: 2.5 ms cathodic + 2.5 ms<br>anodic (DC = 50%)<br>500Hz: 500 us cathodic + 500 us<br>anodic (DC = 50%)<br>1000Hz: 250 us cathodic + 250 us<br>anodic (DC = 50%)<br>5000Hz: 50 us cathodic + 50 us anodic<br>(DC = 50%) | 1, 10, 25, 50,<br>75, 100, 125,<br>150, 175, 200,<br>225, 250, 275,<br>300 μA (peak<br>current) |
| **Asymmetric (ACB 1:4)** | 1:4<br>(cathodic<br>phase 4x<br>amplitude, 1/4<br>duration) | 20, 50, 100,<br>500, 1000,<br>5000 Hz | 20Hz: 5 ms cathodic + 20 ms anodic<br>(DC = 50%)<br>50Hz: 2 ms cathodic + 8 ms anodic (DC<br>= 50%)<br>100Hz: 1 ms cathodic + 4 ms anodic<br>(DC = 50%)<br>500Hz: 200 us cathodic + 800 us<br>anodic (DC = 50%)<br>1000Hz: 100 us cathodic + 400 us<br>anodic (DC = 50%)<br>5000Hz: 20 us cathodic + 80 us anodic<br>(DC = 50%) | 1, 10, 25, 50,<br>75, 100, 125,<br>150, 175, 200,<br>225, 250, 275,<br>300 μA (anodic<br>phase current) |

### Behavioral Testing

In addition to two-photon imaging, each animal underwent behavioral testing to identify the stimulation parameters that would elicit a grimace response in awake rats. The purpose of this test was to determine the overlap between efficacious (RGC activation) and “well-tolerated” or “behaviorally-silent” (no behavioral change or grimace) parameters in awake animals. To achieve this, awake rats underwent EF stimulation as previously described [4]. Two independent observers, blinded to the stimulation parameters administered by a third experimenter, speculated the rat through the open roof of the cage. Both observers consulted the Rat Grimace Scale [20], and, to ensure safety, any response defined as a Grimace Scale greater than or equal to one was defined as the “tolerance” limit. In other words, if the rat exhibited any behavioral change, this amplitude and frequency combination was deemed “intolerable.” Frequencies, amplitudes, and waveforms were systematically swept, maintaining a constant frequency while gradually increasing the amplitude until a grimace or behavioral change was observed. Subsequently, a new frequency was selected, and amplitude sweeping was repeated. The order of frequency selection was varied to avoid data skew arising from potential adaptive or sensitization responses over the duration of the experiment. Tolerance thresholds were mapped as functions of waveform, frequency, and amplitude.

### Two-Photon Imaging Setup

*In vivo* calcium imaging of RGC activity was performed using a transpupillary two-photon microscopy (TPM) system, specifically an upright Olympus FVMPE-RS multiphoton laser-scanning microscope. The animals underwent imaging three days after electrode implantation. Briefly, the rats were anesthetized with a ketamine/xylazine cocktail (5/37.5 mg/kg) and placed under an Olympus two-photon microscope with a Mai Tai Ti: Sapphire femtosecond laser (Spectra-Physics) tuned to 920 nm for GCaMP6f excitation. The pupils were dilated with topical 1% tropicamide and 2.5% phenylephrine to maximize light entry, and the cornea was hydrated with 0.3% hypromellose ophthalmic gel (GenTeal, Alcon Laboratories, Inc., Fort Worth, TX) to retain moisture. The experimental eye was then covered with a glass coverslip to protect it and provide a flat surface for improved image clarity (figure 2a). To image the retina *in vivo* without motion artifacts, the rat head was stabilized in a stereotaxic frame *(SGM-3, Narishige Scientific Instrument Lab, Tokyo, Japan*), laterally rotated, and angled superiorly to align the visual axis with the optical path from the two-photon objective.

The retina was visualized using a Nikon 10X air objective lens with a 16 mm working distance, which offers a wide field of view and provides a greater region of interest (ROI) to capture population-level RGC activity across the retina [21, 22]. A schematic of the imaging pathway is shown in figure 2b. The imaging methodology employed was as follows. First, the retina was brought into focus using the microscopy light imaging mode, and then the two-photon laser was turned on. After entering the RGC layer, the optic disc region was scanned to identify the most robust ROI on the retina for functional fluorescent imaging [23]. A fast resonant scanning mode was utilized to achieve frame rates of 30 Hz for continuous recording, which were sufficient to track calcium transients. The imaging system was carefully calibrated to prevent laser-induced retinal damage. The laser power at the cornea was kept as low as possible to obtain a retinal image. Scanning was limited to 20 min per eye, with three three-minute rest periods between trials, and less than 2.5 minutes of continuous recording in each trial. The emitted fluorescence from GCaMP6f was scanned and detected using photomultiplier tubes with appropriate filters (495-540 nm) to collect calcium-dependent fluorescence signals [24]. During the simultaneous electrical stimulation and calcium imaging experiments, the electrical stimulator was synchronized with the image acquisition process.

**Figure 2:**
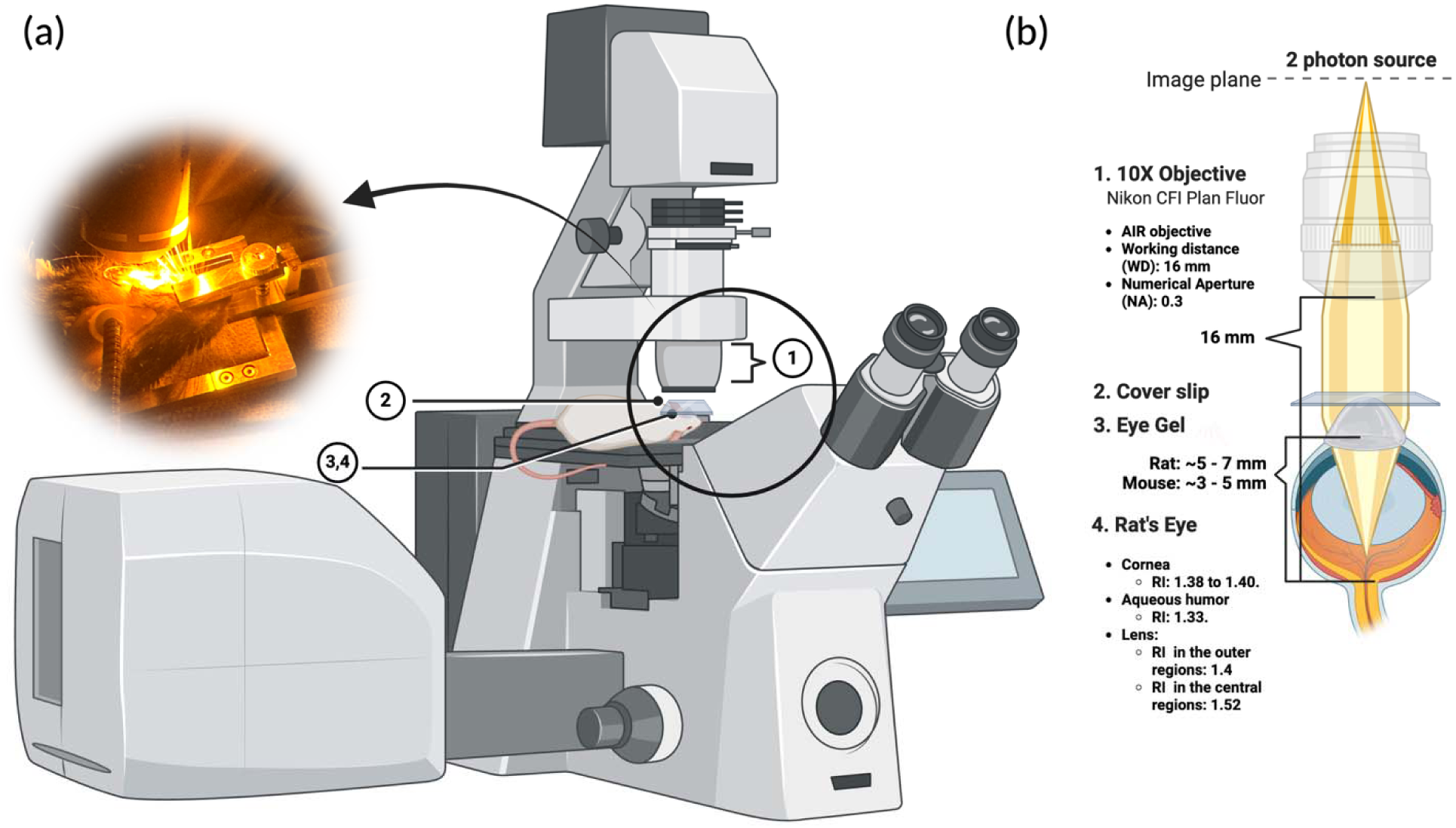
Schematic of imaging setup. (a) Two-photon microscope setup schematic and photograph of an anesthetized rat in the imaging apparatus, illustrating the head-fixed positioning and the objective lens aligned with the eye. (b) Schematic of the optical setup, showing the laser light path through the dilated pupil onto the retina.

### Calcium Activity Extraction and Analysis

For each experimental trial, the laser was turned on for ten seconds without electrical stimulation to establish a resting fluorescence. Then, a designated electrical stimulus (specified by waveform type, frequency, and current amplitude) was delivered for 4 seconds, followed by a 6-second rest period before the next amplitude was tested. Current amplitudes were systematically increased in a stepwise manner, and the stimulation trial was stopped when the stimulus produced a movement artifact precluding the objective’s ability to maintain focus on the same RGC population. This sequence (pre-stim baseline, stimulation, and post-stim) was repeated for multiple conditions in each animal, with at least three minutes of rest between trials to allow for intra- and extracellular calcium concentrations to rebalance and prevent overlap of effects. The order of frequency and waveform type was rotated to avoid systematic bias across rats. Data were acquired from a total of N = 5 rats. All imaging data were saved as time-lapse image stacks (TIFF format) and indexed by stimulus condition for offline analysis.

Calcium imaging data were processed using custom Python scripts available on [<u>Google Drive</u> <u>Link</u>]. Motion correction was not needed because the stereotaxic headframe eliminated breathing artifacts. The fluorescence intensity F(t) over time was calculated by averaging the pixel fluorescence values within the ROI. Baseline fluorescence F₀ was defined for each trial as the mean fluorescence during the one second immediately before the stimulus onset. The fluorescence change relative to baseline was then computed as ΔF/F₀ = (F(t) – F₀) / F₀ for each time point [25]. To quantify RGC activation, two main metrics were calculated from the ΔF/F₀ traces during each stimulation trial: (1) the peak ΔF/F₀ amplitude: the maximum value of the ΔF/F₀ during the stimulation ON time was used as the effectiveness metric of those stimulation parameters; and (2) a binary activation outcome: whether the specific electrical stimulus design showed a “significant” response, defined by ΔF/F₀ exceeding a threshold of 10% increase at any point during stimulation ON time. The 10% threshold was selected based on observed baseline variability, as it consistently exceeded the maximum noise level across all trials. Because baseline fluctuations never surpassed 10%, any normalized fluorescence increase of 10% or more was confidently interpreted as a genuine calcium influx event [26, 27].

### Statistics

Statistical analysis was performed using two-way ANOVA with Tukey’s multiple comparison test to evaluate differences across groups. An alpha level of 0.05 was used, with statistical significance set at p < 0.001. The experimental design involved independent samples, and multiple comparisons were conducted by comparing all groups with appropriate correction applied using Tukey’s method.

### Supplemental Materials

All codes and recordings used in this study are publicly available on [<u>Google Drive Link</u>]. This includes all raw recordings, as well as a sample, labeled, and processed version of the video. To manage output file sizes, most recordings are provided in their raw form, with Python scripts for labeling and generating compressed video versions included. The repository also contains the code used to create all figures and supplementary figures.

## Results

### In Vivo Dose-Dependent RGC Activation with Alternating Current Stimulation

Transpupillary two-photon microscopy (TPM) evaluation of RGC activity *in vivo* revealed a robust RGC response to EF stimulation via extra-ocular electrodes, evident as increases in somatic GCaMP6f fluorescence (ΔF/F₀) coinciding with stimulus delivery. Under baseline (non-stimulated) conditions, fluorescence traces remained stable, reflecting low spontaneous activity. Upon initiation of electrical stimulation, a subset of RGCs rapidly exhibited calcium transients, confirming the effective depolarization and intracellular calcium influx driven by external EF. These effects were reproducible across trials and independent of photic input, as all experiments were performed in the dark to avoid photoreceptor-mediated light stimulation.

To systematically characterize the amplitude-response relationship to EF stimulation, *in vivo* amplitude sweep experiments were conducted at frequencies of 20, 50, 100, 500, and 1,000 Hz. For each frequency, the current amplitude was gradually increased from 1 µA to 300 µA in steps during continuous imaging until a microscopic shift in the head was observed. As this head movement caused the retina to shift out of focus, higher amplitudes were not tested. ACB 1:4 stimulation at 20 Hz demonstrated increasing fluorescence responses with increasing amplitude when the stimulation amplitude exceeded the activation threshold (∼75 µA; figure 3). Stimulation with ACB at 1:4 amplitudes below 50 µA elicited negligible changes in average fluorescence. However, once the current exceeded 75–100 µA, there was a sharp and sustained increase in ΔF/F₀, and response amplitudes scaled proportionally with current intensity. At 150 µA, the calcium response demonstrated the widespread activation of RGCs (figure 3b–3e). The baseline frame (3b) showed minimal fluorescence, whereas stimulation at 50 µA (3c) induced weak, localized activity in a few cells. At 100 µA (3d), a larger population of RGCs exhibited bright GCaMP6f fluorescence, which intensified further at 150 µA (3e). The spatial expansion and intensity of these signals confirmed an apparent dose-dependent effect with enhanced recruitment of RGCs. Together, these data establish that ACB waveform stimulation via an extra-ocular and intracranial electrode can robustly activate RGCs in vivo, with response strength increasing predictably with the stimulation amplitude.

**Figure 3:**
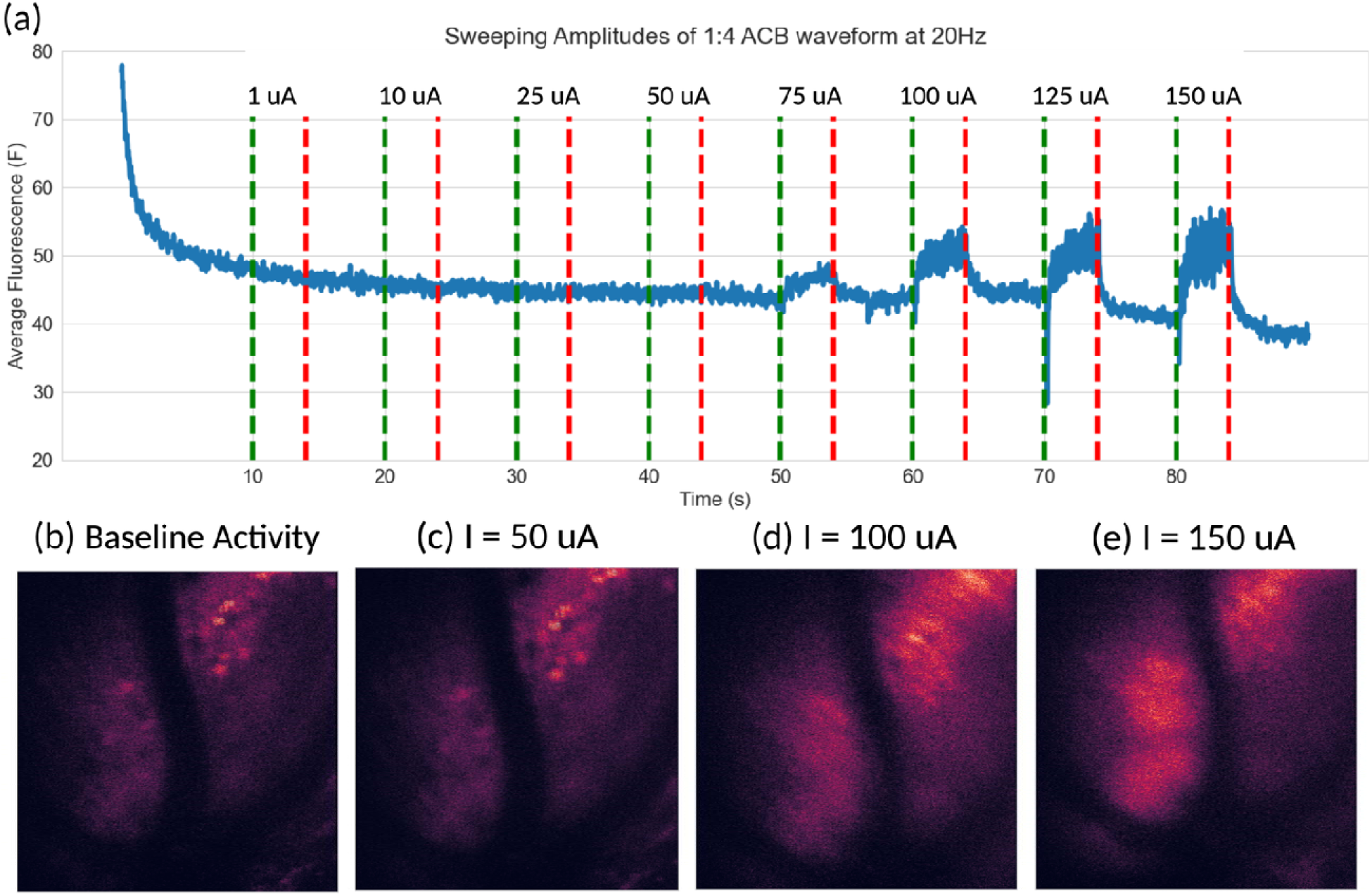
Amplitude sweep of the ACB waveform at 20 Hz demonstrates dose-dependent RGC activation. (a) Average fluorescence intensity (F) from a population of RGCs over time during application of a 1:4 ACB waveform at increasing current amplitudes. Vertical dashed lines mark stimulation epochs: red lines indicate stimulus onset, and blue lines indicate stimulus offset. Fluorescence did not change at low amplitudes (<50 µA) but sharply increased at higher intensities (≥100 µA), indicating robust calcium influx and neural activation. (b–e) Two-photon fluorescence images of the RGC field at baseline and during representative stimulation amplitudes (50, 100, and 150 µA). Increasing fluorescence with brighter activation (yellow-orange) confirmed the amplitude-dependent recruitment of RGCs *in vivo*.

Comparatively, stimulation trains using the SCB 1:1 waveform produced lower response magnitudes and only at higher stimulation amplitudes when compared to ACB 1:4. As shown in figure 4, reliable activating responses, defined as a greater than 10% increase in fluorescence, were only observed at amplitudes exceeding 200 μA for SCB 1:1 across all frequencies tested. In contrast, ACB 1:4 evoked comparable responses at 50–100 μA. Furthermore, SCB 1:1 response occurred only at selected frequencies (20 Hz, 50 Hz, 500 Hz, and 1 kHz), suggesting inconsistent activation across conditions. Even when the responses were elicited, their magnitudes were consistently lower than those achieved with ACB 1:4, despite requiring significantly higher stimulation amplitudes. For instance, at 50 Hz, SCB 1:1 produced a >40% increase in fluorescence only at 300 μA, whereas ACB 1:4 at the same frequency elicited a >50% increase at 75 μA. This highlights the increased activation efficiency of the ACB 1:4 waveform compared to the SCB 1:1.

### Amplitude-Dependent, Frequency-Dependent, and Waveform-Dependent Response Characteristics

Several interesting trends were observed in this study. First, higher current amplitudes led to greater RGC activation across all conditions. This dose-response relationship, shown in figures 4 and 5, was roughly monotonic: at 20 Hz, the normalized score increased from 0.14 at 50 µA up to 0.85 at 150 µA. Second, lower stimulation frequencies generally resulted in stronger calcium responses than those at higher frequencies. For instance, at stimulation amplitudes of 100–150 µA, 20 Hz, and 50 Hz, the stimulations elicited notably larger and more widespread calcium transients than responses observed at 500 Hz or 1 kHz. The increased response at 20 Hz was significantly higher than that at frequencies of 100 Hz or greater (two-way ANOVA with Tukey’s multiple comparison test, p < 0.0001). Finally, the asymmetric 1:4 waveform outperformed the symmetric 1:1 waveform. In figure 4a, the cells corresponding to ACB conditions were consistently warmer in color, indicating greater fluorescence changes than their SCB counterparts in the same column. At 20 Hz, asymmetric waveforms produced significantly greater calcium responses than did symmetric waveforms, highlighting their superior efficacy (p < 0.001; ANOVA). Specifically, stimulation with the 20 Hz ACB 1:4 condition elicited significantly greater responses compared to all SCB 1:1 frequencies and amplitudes tested. However, there was no significant difference between the 1:4 ACB and 1:1 SCB waveform outcomes at higher frequencies, highlighting the decreased effectiveness of high-frequency stimulation for RGC activation.

The probability of activation, or the fraction of rats showing a ΔF/F₀ greater than 10%, complements the amplitude heatmap (figure 4b). This is important because it demonstrates that, despite inter-rat variability, RGC activation can be reliably achieved if the waveform is asymmetric and the amplitude exceeds cellular thresholds. In terms of consistency, the waveform effect was again evident: the ACB waveform (1:4) consistently yielded higher activation probabilities than the SCB (1:1) waveform. Under certain conditions, the difference was striking; for example, at 50 Hz and 150 µA, the score was 1, indicating that ACB 1:4 elicited calcium transients in all the rats tested. Comparatively, SCB 1:1 at the same frequency, with an amplitude of at least 200 µA, was required to achieve the same probability of activating RGCs (>10% ΔF/F₀). This suggests that stimulation with an asymmetric pulse not only increases the magnitude of the response in individual neurons but also engages a greater fraction of the RGC population at any given stimulus strength. From a practical standpoint, the 1:4 waveform can therefore achieve a higher desired level of network activation at a lower current intensity than the 1:1 waveform, potentially offering efficiency benefits for therapeutic stimulation.

**Figure 4:**
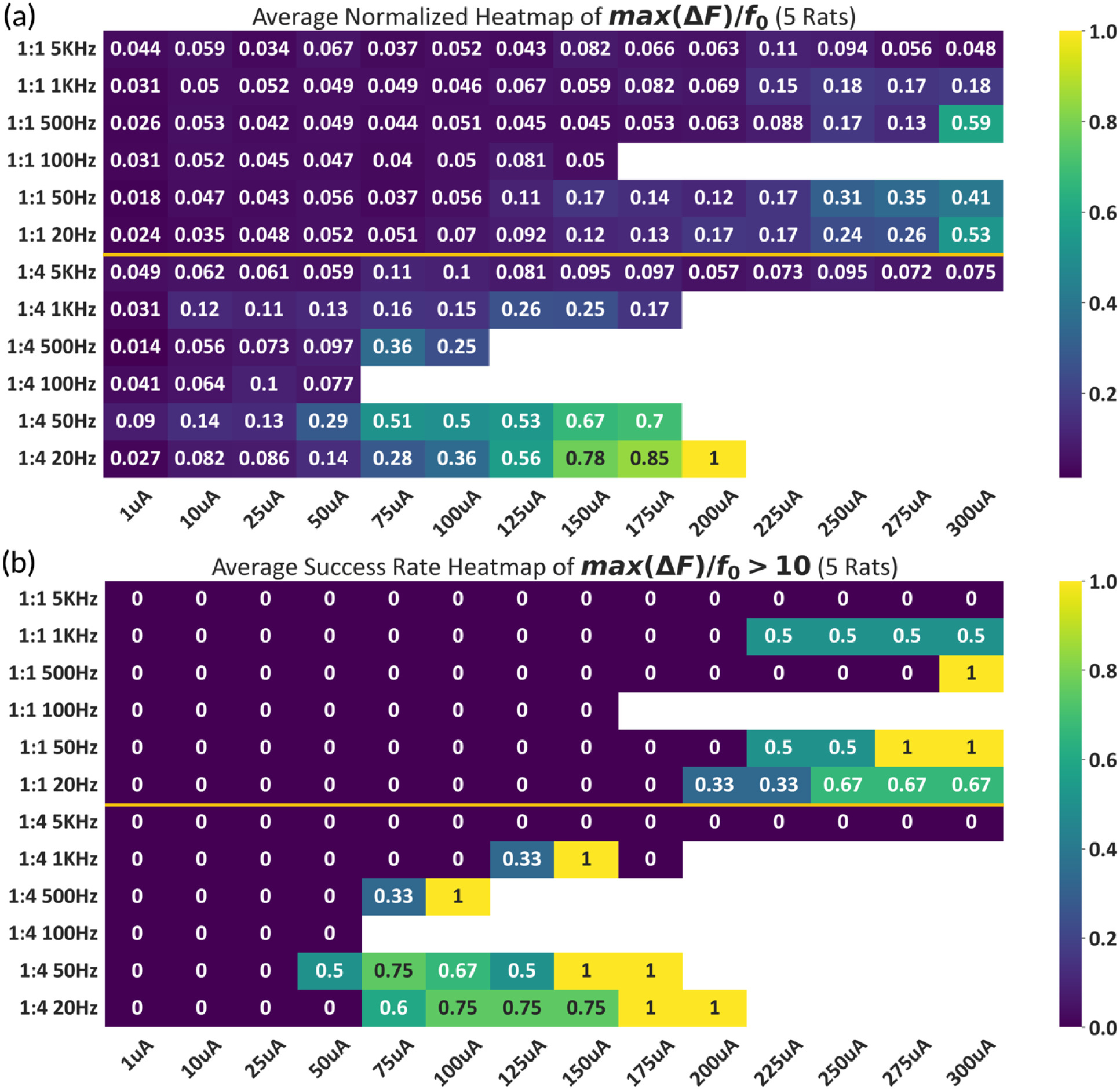
Comparative analysis of stimulation efficacy using SCB 1:1 and ACB 1:4 waveforms across frequencies and amplitudes. (a) Heatmap of the average normalized peak calcium response (ΔF/F₀) measured in RGCs across five animals. Rows indicate different waveform configurations and stimulation frequencies, whereas columns represent increasing current amplitudes (1–300 µA). Brighter colors (yellow-green) indicate stronger calcium responses. The asymmetric charge-balanced (ACB, 1:4) waveform consistently produced higher calcium responses, particularly at lower frequencies (20 and 50 Hz). (b) Corresponding heatmap of the average success rate, defined as the fractio of trials in which the calcium response exceeded the threshold (ΔF/F > 0.10). Success rates increased significantly with higher amplitudes and were markedly higher for the 1:4 waveform at frequencies of 20 Hz and 50 Hz. Together, these heatmaps illustrate the superior effectiveness of the 1:4 waveform configuration in eliciting robust RGC activation across the tested stimulation parameters.

**Figure 5:**
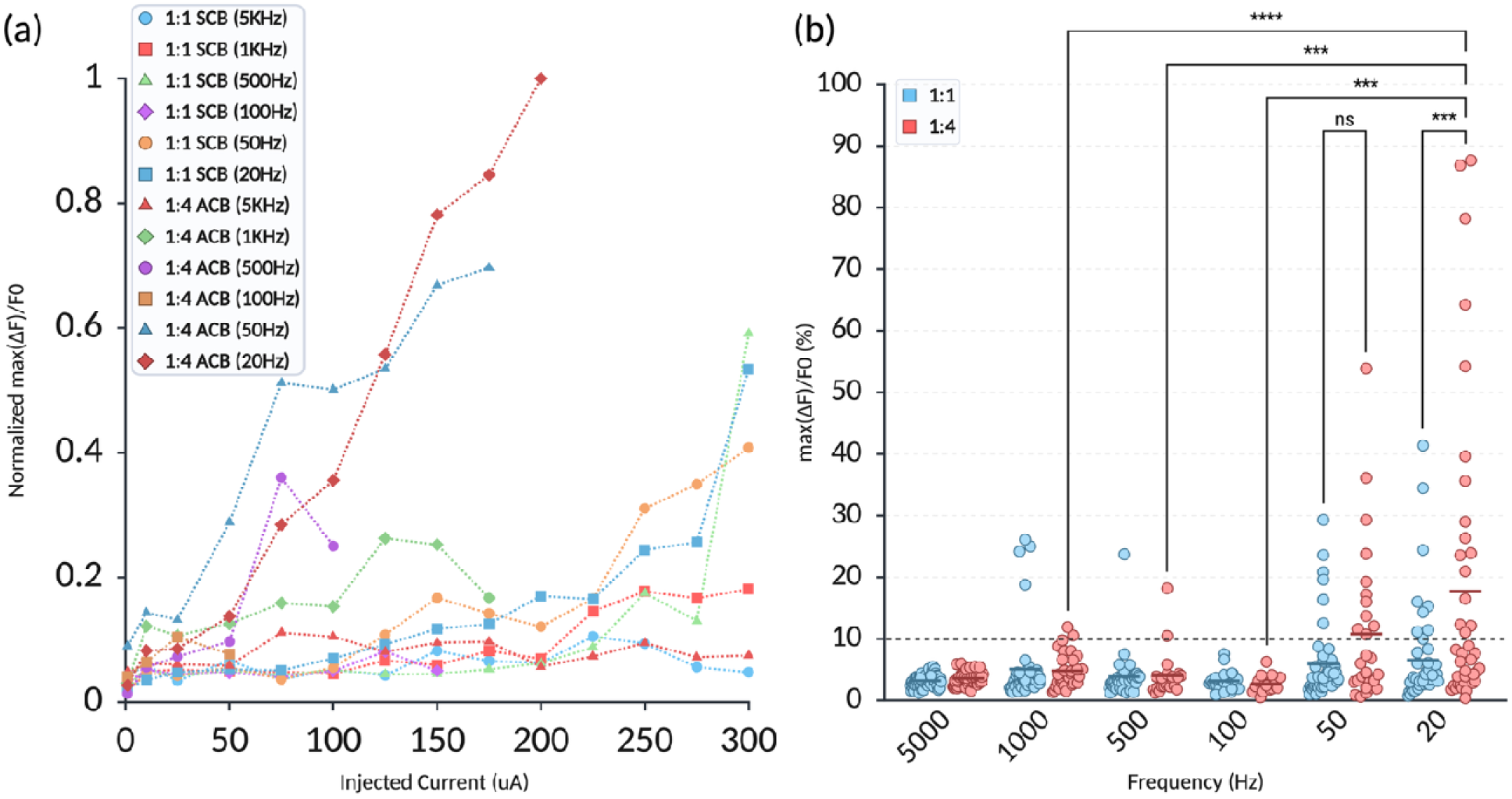
Comparison of RGC calcium responses to SCB and asymmetric ACB waveforms across varying frequencies and current amplitudes. (a) Normalized peak calcium responses (ΔF/F₀) plotted against injected current amplitudes (1–300 µA) for different waveform-frequency combinations. Notably, the ACB 1:4 waveform exhibits greater normalized responses at lower current intensities, especially at 20 Hz and 50 Hz. (b) Statistical comparison of peak calcium responses across frequencies between SCB (1:1, blue circles) and ACB (1:4, red circles) waveforms. Each dot represents the results of individual trials pooled across multiple animals. Horizontal lines represent the group means. Significance is indicated by asterisks (*** P <0.0001, ns: not significant). Asymmetric waveforms at lower frequencies (20 Hz, 5 Hz) elicited significantly greater peak calcium responses than higher frequencies and symmetric waveforms.

### Behavioral responses to electrical stimulation of the optic nerve

To assess the response to electrical stimulation of the optic nerve in awake, freely moving animals with implanted electrodes, a series of behavioral tests was conducted across different frequencies for both waveform types. Tolerance threshold was defined as the maximum current amplitude that could be delivered without inducing a physical response (e.g., blinking, twitching, startle, excessive tearing, altered respiration, or further behavioral alterations). A shown in figure 6, animal tolerance is proportional to pulse duration, with animals tolerating stimulation with higher amplitudes when the pulse duration is limited.

**Figure 6:**
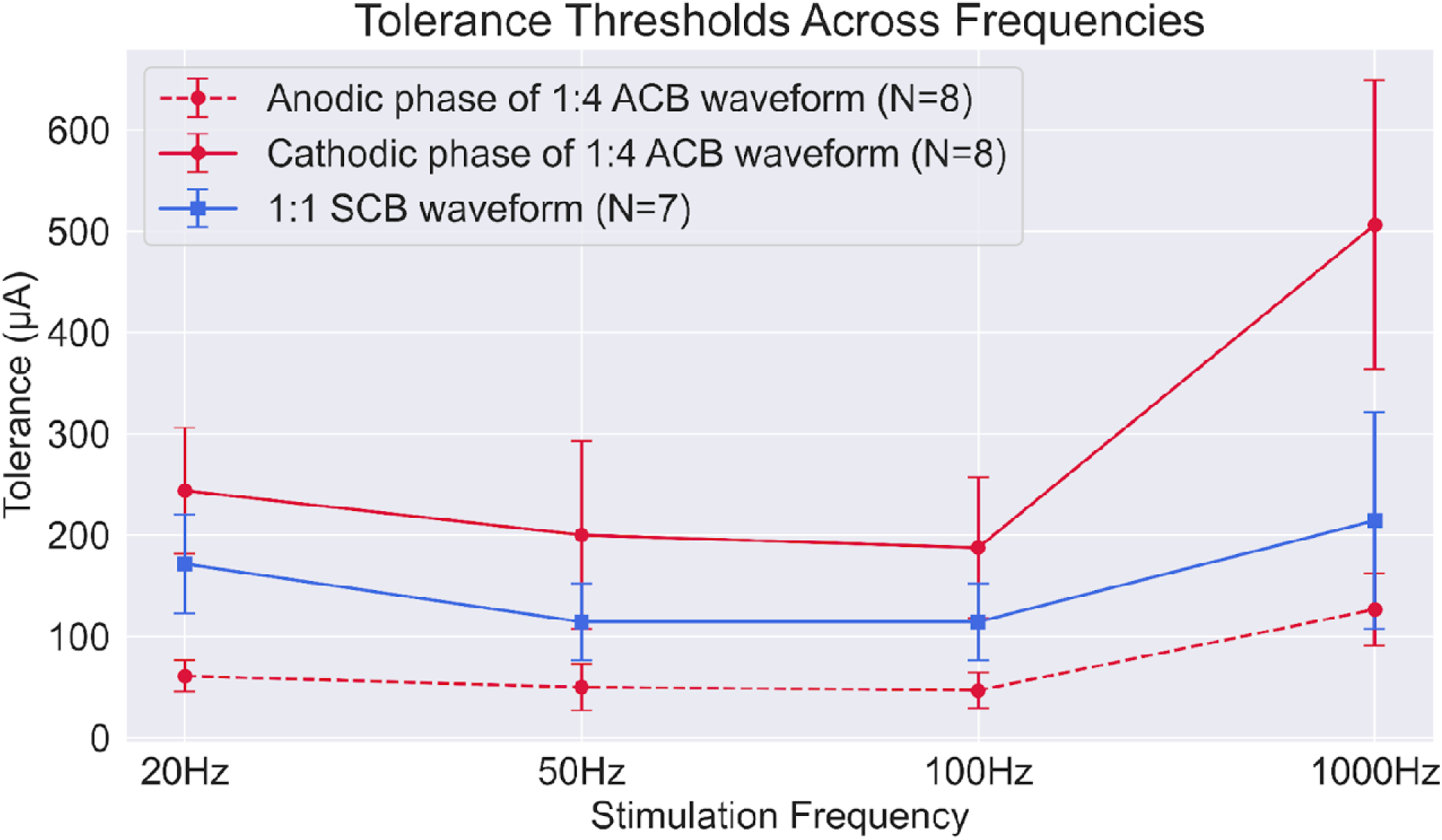
Maximum tolerable stimulation amplitudes across frequencies for 1:1 and 1:4 waveform configurations. The tolerance threshold (defined as the maximum current amplitude tolerated without any behavioral response) was plotted across multiple frequencies (20 Hz, 50 Hz, 100 Hz, 1000 Hz). Error bars represent standard deviation across animals.

### Therapeutic Window Analysis: Superior Operation Range of ACB 1:4 Waveform Compared to SCB 1:1

*In vivo* activation and tolerance thresholds were systematically compared across multiple frequencies (20, 50, 100, 500, and 1000 Hz) for asymmetric (1:4) and symmetric (1:1) waveforms to evaluate the practical and safe therapeutic windows for retinal neuromodulation in awake, alert subjects. Activation thresholds were defined as the minimum amplitude at which RGCs reliably exhibited measurable calcium responses (ΔF/F₀ > 10%). In contrast, tolerance thresholds are the maximum current amplitude that can be applied without eliciting a noticeable behavioral reaction. The symmetric (1:1) waveform exhibited no practical therapeutic window across all frequencies tested, as there was no overlap between the waveform amplitudes that elicited RGC responses and those tolerated in awake animals (Figure 7). In contrast, the asymmetric (1:4) waveform demonstrates a close alignment between the activating parameters and tolerance levels across frequencies. This overlap is particularly apparent at 500 Hz, where the activation amplitudes approach the tolerance threshold.

These observations were supported by a one-sided Mann–Whitney U test. This test evaluated whether the activation thresholds were significantly greater than the tolerance thresholds. For the symmetric (1:1) waveform, the test yielded a statistically significant result (p = 0.012), confirming that the activation thresholds exceeded tolerance and indicated a lack of an overlapping amplitude range. In contrast, the 1:4 waveform at 50 Hz did not show a significant difference between activation and tolerance thresholds (p = 0.14), suggesting that asymmetric stimulation offers a closer alignment between efficacy and safety, potentially enabling a more viable stimulation range.

**Figure 7:**
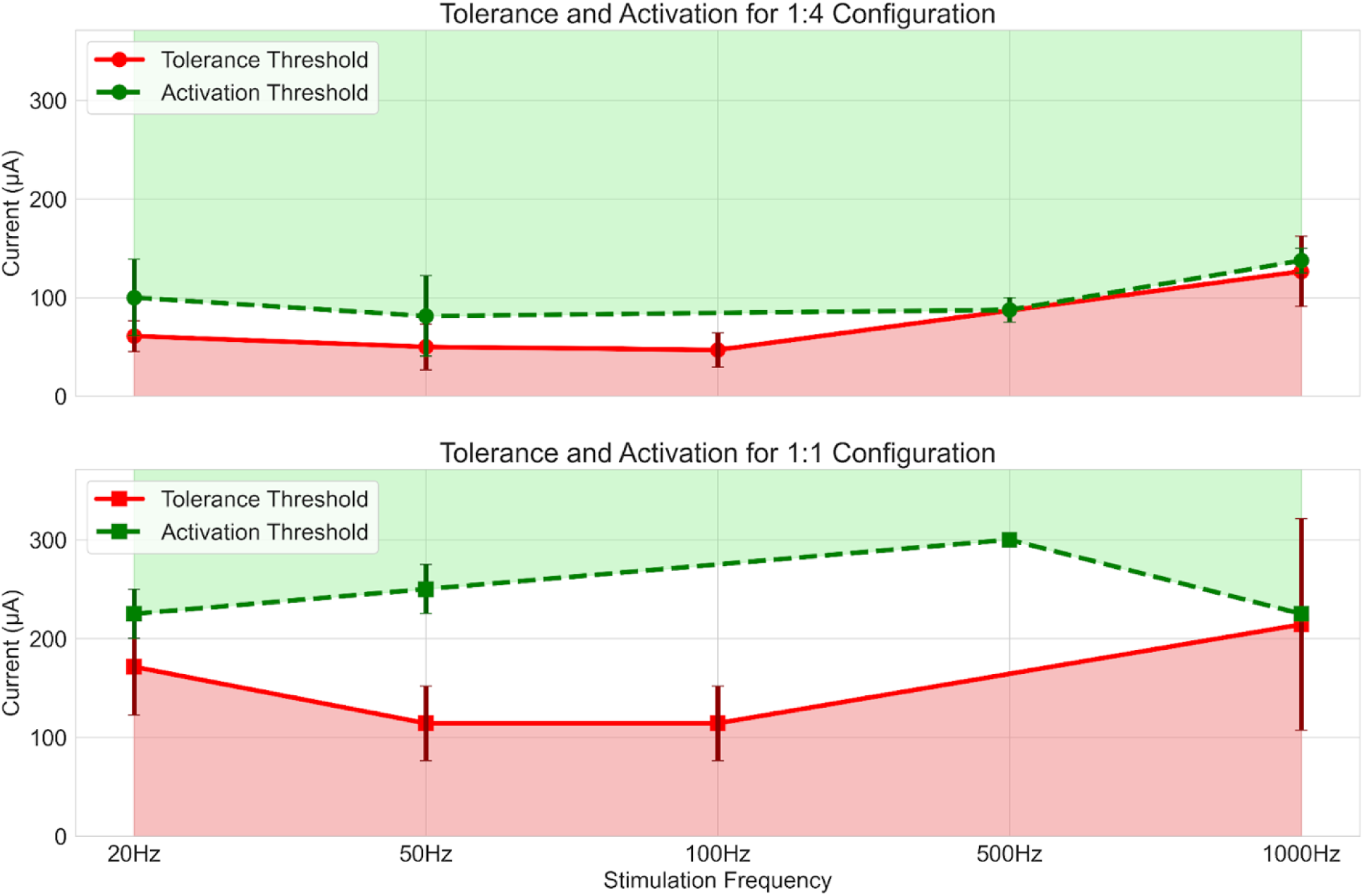
Comparison of activation and tolerance thresholds between the ACB 1:4 and SCB 1:1 waveform configuration. The ACB 1:4 waveform consistently exhibited lower activation and tolerance thresholds across the tested frequencies. Notably, despite these lower amplitude limits, the ACB 1:4 configuration maintained a measurable therapeutic window at several frequencies, defined as the range between the minimum anodic amplitude required to evoke reliable RGC activation (ΔF/F₀ > 10%) and the maximum amplitude tolerated without inducing behavioral responses. In contrast, the SCB 1:1 waveform demonstrated a higher absolute anodic tolerance threshold but an even higher activation threshold, resulting in no observable therapeutic window, particularly at mid-rang frequencies.

## Discussion

Using transpupillary two-photon calcium imaging to validate optic nerve stimulation parameters in vivo, this study showed that EF stimulation from extra-ocular electrodes can directly activate RGC somas. We further show that ACB stimulation consistently produces stronger and more reliable activation than SCB stimulation. Additionally, behavioral experiments indicate that ACB 1:4 provides a therapeutic window not observed with SCB waveforms. Among the tested parameters, the ACB 1:4 waveform at 20–50 Hz produced the most consistent and robust response. To our knowledge, this is the first direct in vivo demonstration that RGCs can be modulated by electric fields delivered through extraocular electrodes placed within the orbit and cranium, establishing a less invasive alternative to intraocular electrode placement and providing a foundation for neuromodulation strategies targeting the visual pathway [28–35]. For certain indications, extraocular electrodes offer practical advantages including safer implantation and improved long-term accessibility compared to epiretinal and subretinal neuroprostheses.

The observed frequency- and amplitude-dependent modulation of RGC activity is consistent with prior reports emphasizing the critical roles of waveform shape and stimulation frequency in defining neural activation thresholds. As previously shown [19, 34], moderate stimulation frequencies (20–50 Hz) elicited stronger calcium responses than higher frequencies (500–1000 Hz). This difference may reflect reduced neuronal excitability or calcium saturation at higher frequencies [36–38], possibly because of insufficient recovery time between stimuli. Similar patterns of frequency-dependent activation of RGCs have been reported in previous retinal studies. For example, Freeman et al. demonstrated layer-specific responses to stimulation frequency, with 5–10 Hz primarily activating photoreceptors, 25 Hz targeting bipolar cells, and 100 Hz selectively engaging RGCs [35]. Our in vivo findings extend the ex vivo observations by confirming that recovery-dependent frequency tuning governs RGC excitability in the intact visual pathway. Importantly, unlike prior in vitro work, these results demonstrate that low-frequency ACB stimulation delivered via extraocular electrodes can modulate or activate RGCs in living animals while remaining tolerable in awake subjects.

The superior efficacy of the 1:4 ACB waveform compared with the 1:1 SCB waveform likely arises from the physiological consequences of waveform asymmetry and rebound excitation. One possible mechanism is that initiating the pulse with a brief, high-amplitude cathodic current provides a transient hyperpolarizing effect, which facilitates stronger excitation when the waveform transitions to a longer, lower-amplitude, anodic phase that depolarizes tissue near the target [1]. This sequence more effective drives the membrane potential toward depolarization threshold, promoting activation of voltage-gated calcium channels [39]. Prior work support this interpretation: asymmetric biphasic pulses can reduce activation thresholds by up to 55% compared to symmetric pulses [40]; anodic-first configurations in the Argus II trial lowered RGC thresholds in human participants [39]; and in the rabbit optic nerve, short cathodic pulses elicited larger responses with lower current requirements than alternative waveforms [41]. Frequency-dependent decreases in electrically evoked potentials have also been consistently reported across species [42–44]. Together with our findings, these observations indicate that waveform asymmetry and frequency spacing act synergistically to lower the thresholds and enhance RGC recruitment in vivo.

The translational relevance of these findings lies in the presence of a therapeutic window for the 1:4 ACB waveform characterized by both lower activation thresholds and improved behavioral tolerance. Increased waveform asymmetry and shorter cathodic duration may reduce the net charge accumulation, limit electrode degradation, and improve subject comfort, all of which are critical considerations for long-term use and clinical translation. By shifting the effective stimulation range into the tolerable amplitude window, the 1:4 ACB waveform emerges as a potential candidate for vision restoration strategies targeting amblyopia, glaucoma, and optic neuropathies, whereas SCB stimulation often required amplitudes exceeding tolerance before producing reliable activation. The ability of low-frequency ACB stimulation to reliably drive calcium influx further suggests applications in neuromodulation therapies aimed at promoting plasticity, synaptic strengthening, and neuroprotection [45–49]. Accordingly, the 1:4 ACB waveform represents both a technical advance in RGC stimulation and a practical step toward clinically viable neuromodulation therapies.

One possible explanation for the near-perfect alignment between the 1:4 ACB activation threshold and tolerance threshold amplitude (Figure 7) is that animals may exhibit a startle response to visually evoked percept rather than discomfort from the stimulus itself. Direct optic nerve stimulation likely generates visual sensations such as flashing lights or “photopsia,” analogous to the phosphene response commonly experienced by patients undergoing rTMS of the visual cortex [50–52]. In contrast, for 1:1 SCB waveform, tolerance thresholds were typically below activation thresholds across tested frequencies, suggesting that behavioral responses were more likely due to discomfort than a visually driven startle response.

This study had several limitations. Calcium imaging is an indirect measure of neuronal activity and may underestimate total spiking [53–56]. Because stimulation was delivered using an extraocular electrode paired with an intracranial return, electric fields were likely strongest at the axons rather than somas. Given the limited ability of axonal action potentials to back-propagate to the soma, the responses detected here may underestimate the degree of RGC activation [57, 58]. Additionally, the use of large-field stimulation prevented the assessment of spatial selectivity with respect to electrode size and placement. These factors suggest that the observed effects likely represent a conservative estimate of ACB efficacy, underscoring the need for targeted electrode designs in future studies.

Future investigations should evaluate long-term safety under repeated stimulation, refine electrode designs to improve spatial precision besides moving the return electrode to the scalp, and incorporating behavioral assays capable of distinguishing therapeutic and adverse responses. Extending this approach to disease models will be essential to establish clinical efficacy, particularly by correlating neural activation with functional visual outcomes, such as contrast sensitivity, optokinetic responses, and visual evoked potentials. Ultimately, linking cellular-level activation to measurable improvements in vision will be a decisive step in translating these findings to human therapy.

## Conclusion

This study provides direct evidence that extra-ocular electrode stimulation can activate RGCs in vivo. We identify the 1:4 asymmetric charge-balanced (ACB) waveform as an efficient and well-tolerated paradigm for driving RGC activation that is more robust and reliable than a symmetric 1:1 waveform under equivalent conditions, remaining better tolerated in awake animals. Delivery of EFs via extra-ocular electrodes offers advantages in electrode safety and removal compared to intra-ocular approaches, an increasingly important consideration for chronic neuromodulation systems. By combining waveform asymmetry with optimal frequency spacing, this strategy enables stronger activation at lower thresholds, while maintaining a therapeutic window compatible with clinical translation. Together, these findings strengthen the rationale for ACB waveforms use in therapeutic neural stimulation, refine mechanistic understanding, and establish a practical framework for developing targeted, safe, and effective EF-based neuromodulation therapies for vision restoration.

## Acknowledgements

NH was supported by a Research to Prevent Blindness (RPB) Medical Student Research Fellowship grant. KKG was supported by grants from the NEI/NIH (R01EY035375) and NSF (2121164). MSB was supported by grants from the NIA/NIH (K01AG066847) and NSF (2121164). GL and OS were supported by an NSF grant (2121164). This work was also supported by an unrestricted grant to the Department of Ophthalmology at the University of California Irvine and University of Southern California from Research to Prevent Blindness and NEI (P30EY029220, P30 EY034070). The content is the sole responsibility of the authors and does not necessarily represent the official views of the National Institutes of Health. No funding sponsors were involved in the study design, collection, analysis, interpretation of data, writing of the report, or the decision to submit the article for publication.

## Ethical Statement

All animal procedures complied with the ARRIVE guidelines and the ARVO Statement for the Use of Animals in Ophthalmic and Vision Research and were approved by the Institutional Animal Care and Use Committee at the University of Southern California [16].

